# Retinal specializations to a micro-predatory and crypto-benthic life-style in the Mediterranean triplefin blenny *Tripterygion delaisi*

**DOI:** 10.1101/208298

**Authors:** Roland Fritsch, Shaun P. Collin, Nico K. Michiels

**Affiliations:** Animal Evolutionary Ecology, Institute of Evolution and Ecology, Department of Biology, University of Tübingen, Tübingen, Germany.; The Oceans Institute and the School of Biological Sciences, The University of Western Australia, Crawley, Western Australia, Australia.

**Keywords:** *Tripterygion delaisi*, visual ecology, retinal wholemount technique, retinal specializations, retinal topography, visual acuity, cone mosaic, triple cones

## Abstract

The environment and lifestyle of a species are known to exert selective pressure on the visual system, often demonstrating a tight link between visual morphology and ecology. Many studies have predicted the visual requirements of a species by examining the anatomical features of the eye. However, among the vast number of studies on visual specializations in aquatic animals, only a few have focused on small benthic fishes that occupy a heterogeneous and spatially complex visual environment. This study investigates the general retinal anatomy including the topography of both the photoreceptor and ganglion cell populations and estimates the spatial resolving power of the eye of the Mediterranean triplefin *Tripterygion delaisi*. Retinal wholemounts were prepared to systematically and quantitatively analyze photoreceptor and retinal ganglion cell densities using design-based stereology. To further examine the retinal structure, we also used magnetic resonance imaging and histological examination of retinal cross sections. Observations of the triplefin's eyes revealed them to be highly mobile, allowing them to view the surroundings without body movements. A rostral aphakic gap and the elliptical shape of the eye extend its visual field rostrally and allow for a rostro-caudal accommodatory axis, enabling this species to focus on prey at close range. Single and twin cones dominate the retina and are consistently arranged in one of two regular patterns, which may enhance motion detection and color vision. The retina features a prominent, dorso-temporal, convexiclivate fovea with an average density of 104,400 double and 30,800 single cones per mm^2^, and 81,000 retinal ganglion cells per mm^2^. Based on photoreceptor spacing, spatial resolving power was calculated to be between 6.7 and 9.0 cycles per degree. Location and resolving power of the fovea would benefit the detection and identification of small prey in the lower frontal region of the visual field.

## Introduction

Vertebrate eyes typically show remarkable specializations to view their surroundings with their ocular anatomy providing a strong indication of each species’ ecology and evolutionary history. While all vertebrate eyes share the same general organization, many species have evolved individual specializations that enhance their visual perception of the environment. These visual specializations often reflect ecological niches, which is particularly true for fishes as they occupy a diverse range of microenvironments. As the largest group of vertebrates, they have evolved diverse lifestyles in a wide range of habitats and these are often reflected in a variety of visual adaptations. The resultant link between a species’ visual ecology and eye anatomy allows one to make meaningful predictions from one about the other (Hughes, 1977). Such investigations are especially informative when little is known about the ecology and behavior of a species, for instance when behavioral observations are difficult.

Adaptive modifications of the basic eye structure to match specific visual demands can be found at all structural levels, e.g. the tubular eyes of some deep sea teleosts (Collin et al., 1997; Warrant and Locket, 2004; Biagioni et al., 2016), the bipartite pupil and asymmetrical lens of the four-eyed fish (Nicol and Somiya, 1989), or multifocal lenses (Kroger et al., 1999). However, the ultimate level of visual perception is the retina, where a range of specializations have been described including variations in photoreceptor number, density, arrangement, type and spectral sensitivity (Collin and Shand, 2003), and the presence and location of a number of types of ocular reflectors (Fritsch et al., 2017). Depending on visual demands, specializations often involve a trade-off between sensitivity and resolution. For instance, high summation rates, grouped receptors, and large receptor diameters can all serve to increase sensitivity, but inevitably lower spatial resolving power (Land and Nilsson, 2012). As a compromise, many species possess distinct retinal areas of high receptor density to mediate higher spatial resolution. Different types and combinations of such areas, varying in shape, size, and retinal position, have been described for many fish species (Collin, 2008).

Which type of specialization occurs in a species has been proposed to correlate with their behavioral ecology and visual environment (Hughes, 1977). A horizontal streak of high receptor density is generally found in species whose environments feature a prominent horizon. This requires a fixed (and often wide) region of high visual acuity across the retina, which enables the eye to scan a large panoramic visual field for predators without the need for pronounced eye movements (Collin and Collin, 1999). In contrast, species that search for cryptic prey in highly structured, heterogeneous environments would be expected to feature a pronounced *area centralis*, a concentric region of increased cell density (Collin and Shand, 2003). In many species with particularly high spatial resolution, the neuronal elements other than the photoreceptors are partially or completely displaced from the center of the *area centralis*. This creates an indentation in the retinal structure, i.e. a fovea, which is typical for birds (Tucker, 2000; Coimbra et al., 2006; Coimbra et al., 2009; Coimbra et al., 2014) and primates (Williams and Coletta, 1987; Wassle et al., 1990; Wikler and Rakic, 1990), but has also been found in many fish species (Collin and Collin, 1988c; 1999; Collin et al., 2000; Collin and Shand, 2003). Blennioid species, which usually live in complex 3D environments, have also been known to possess foveae for a long time (Kahmann, 1936). These early studies, however, constituted only macroscopic and qualitative assessments. Since then, only few studies of retinal morphology, visual specializations, and spatial resolving power have been carried out in small, crypto-benthic fish species. Examples include the blenniid *Petroscirtes variabilis* (referred to as *Dasson variabilis* in the original publication), the tripterygiid *Forsterygion varium*, the sandlance *Limnichthyes fasciatus*, and several syngnathid species (Collin and Collin, 1988c; Collin and Pettigrew, 1989; Pankhurst et al., 1993; Collin and Collin, 1999; Lee and O'Brien, 2011). Even these more recent studies, however, often lack a systematic analysis of the retinal topography, or only consider retinal ganglion cells.

Here, we present a comprehensive study that includes the ocular anatomy, the quantitative, topographic distribution of both photoreceptor cells and retinal ganglion cells, and an estimate of the spatial resolving power of the eyes of the Mediterranean subspecies of the yellow black-faced triplefin blenny *Tripterygion delaisi* (Tripterygiidae). *T. delaisi* reaches a length of 3 - 6 cm (SL), occurs on algae-encrusted, rocky substrates at depths of 3 - 40 m in the Mediterranean and Northeast Atlantic, and feeds as an opportunistic micro-predator on a variety of small (1 - 3 mm) invertebrates, especially crustaceans (Wirtz, 1978; Zander, 1982; Zander et al., 1986). Both sexes feature the same mottled, white and brown coloration with darker stripes and bright spots (Wirtz, 1978). This matches their benthic habitat characterized by rocks overgrown with diverse species of algae both in shaded and exposed sites (La Mesa et al., 2005). *T. delaisi*’s behavior contributes to its camouflage, as triplefins spend much of their time perching motionlessly on the substrate, while scanning the environment for potential threats and prey with frequent and extensive eye movements (unpublished observations). Occasionally, *T. delaisi* changes its position with short, impulsive darts, just to freeze again in the new location. Exceptions to its generally cryptic coloration are the males' black-and-yellow breeding coloration (Wirtz, 1978), and the red-fluorescing irides of both sexes whose brightness can be physiologically regulated via melanophores (Wucherer and Michiels, 2014). The general intensity of the fluorescence increases with the depth at which an individual lives (Meadows et al., 2014) in response to the reduced overall light intensity at greater depth (Harant et al., 2016). Furthermore, *T. delaisi*'s set of cone receptors is sensitive to the long-wavelength light produced by its fluorescent pigments and the intensity of the fluorescence is sufficient to create perceivable contrasts (Kalb et al., 2015; Bitton et al., 2017). The function of a fluorescent iris and other unusual external features of the eye of *T. delaisi* and their effect on vision are still unclear. Their potential role is discussed and investigated in detail elsewhere (Fritsch et al., 2017; Harant and Michiels, 2017).

The intricate patterns and changes in coloration, the unusual visual features of its eyes, as well as its frequent eye movements to scan its environment, suggest a rich visual ecology and complex visual system. In this study, we use retinal wholemounts combined with design-based stereology, retinal cross sections, and magnetic resonance imaging to characterize ocular and retinal specializations for *T. delaisi*'s particular micro-predatory, cryptic lifestyle.

## Methods

### Specimens

We used individuals of *T. delaisi* (n = 10) to either prepare retinal wholemounts (4), paraffin-embedded thick histological sections (4), or magnetic resonance imaging scans (2). Individuals used for retinal wholemounts were wild-caught at STARESO research station (Pointe Revellata, BP33 20260 Calvi, Corsica, France) in July 2014. All others specimens were wild-caught at the *Centro Marino Elba* research station (Loc. Fetovaia 72, I-57034 Campo nell'Elba, Italy) in June 2013. All animals were euthanized according to German animal ethics legislation under notifications *AZ. 13.06.2013* and *AZ. 29.10.2014* issued to NKM by the animal welfare department of the district administration of Tübingen ("Regierungspräsidium Tübingen"). Specifically, the fish were immersed in seawater containing a lethal dose of 500 to 1000 mg/l tricaine methanesulfonate (MS-222), adjusted to pH = 8.2 with NaOH, until there was no discernable opercular movement for at least one minute. Cutting the spinal cord with a scalpel ensured euthanasia. We then measured the standard length (SL) of each individual. The eyes were excised from the skull and the horizontal and vertical diameters of the eye and pupil of each fish were measured using digital calipers (to the nearest 0.1 mm).

### Eye mobility

*T. delaisi'*s pronounced eye movements have already been mentioned in the most comprehensive study on the species' behavior (Wirtz, 1978) but no study ever quantified that mobility. We extracted this information from video footage from previous, unpublished work. In these videos *T. delaisi* is seen from above while subjected to an optokinetic reflex experiment, demonstrating its horizontal eye mobility. We scanned the videos of three individuals for the frames with the maximal deflection of the eyes in relation to the midline of the head, exported the frames as still images, and measured the maximal angular change in ImageJ v.1.48t. We are aware that this approach only yields an approximate and incomplete estimate of eye mobility, but these data are nevertheless a meaningful addition to our main results. The implementation of a more exhaustive assessment of eye movements goes beyond our scope to describe this species’ retinal anatomy.

### Histological serial sections and magnetic resonance imaging (MRI)

The four *T. delaisi* prepared for paraffin-embedded thick sections and the two individuals used for magnetic resonance imaging (MRI) are the same as in a previous study (Fritsch et al., 2017), which provides a comprehensive description of the procedures. Briefly, the eyes used for histological sections were immersion fixed with 4 % paraformaldehyde in 0.155 M Sorensen‘s phosphate buffer, and the tissue decalcified, dehydrated and embedded in paraffin blocks. Blocks were sectioned at 10 μm thickness and along all three anatomical planes across the four prepared specimens. The sections were collected on standard microscope slides, stained with phosphotungstic acid haematoxylin (Mallory, 1900), and then mounted with Entellan^®^. Sections from the region of interest were digitized and measurements were acquired in ImageJ 1.48t (Fiji distribution package).

For the MRI scans performed on two specimens, the fish were euthanized as above, and immersion fixed in 4 % paraformaldehyde in phosphate buffered saline, which also contained 0.25 % of a 1 M contrasting agent (Gadovist^®^). Scans were run on a 16.4 T Ultrashield^TM^ Plus 700 WB Avance nuclear magnetic resonance spectrometer with ParaVision^®^ 5.1 software (Bruker BioSpin GmbH, 76287 Rheinstetten, Germany). The digital 3D segmentations of the MRI data were created in ITK-SNAP, version 2.4.0 (Yushkevich et al., 2006).

### Retinal wholemount preparation

Immediately after euthanasia, the four specimens used for retinal wholemount preparation were transferred to modified Ringer's solution (MRS) optimized for marine teleosts, which contained 125.3 mmol NaCl, 2.7 mmol KCl, 1.8 mmol CaCl_2_, 1.8 mmol MgCl_2_, 5.6 mmol D(+)-Glucose, and 5.0 mmol Tris-HCl per liter, buffered to pH = 7.2 (Wucherer and Michiels, 2012). Dissection of the eyes and preparation of the retinal wholemounts are only summarized here, as we largely followed procedures comprehensively described in Coimbra et al. (2006) and Ullmann et al. (2012). We excised each eye from the skull, transferred it to MRS, and removed the dermal cornea, scleral cornea, lens, iris, and vitreous body. Lens diameter was measured to the nearest 0.01 mm with digital calipers. The remaining eyecup was then fixed with 4 % paraformaldehyde in 0.1 M, pH = 7.3, Sorensen's phosphate buffer (SPB) for 20 - 30 min at room temperature just until the previously clear retina started to become translucent. Excess fixative was subsequently removed by washing the eyecup in the buffer three times for one minute each. In all following steps, the tissue was constantly submerged in or moisturized with SPB. We then applied four to five radial cuts to the eyecup that enabled us to consecutively separate the other tissue layers (choroid and retinal pigment epithelium) from the retina, allowing the isolated retina to be flattened onto a non-subbed glass slide. Since we wanted to analyze both photoreceptor and retinal ganglion cell (RGC) populations in the same retinae, and the RGC preparation involves irreversible procedures, the preparation, viewing and analysis of the topography of the photoreceptors was carried out first, as follows.

To obtain a clear view of the photoreceptor layer without damaging it, only loose retinal pigment epithelium (RPE) was removed mechanically, while any remaining pigment was bleached by immersing the retina in a mixture of 5 parts double-concentration phosphate buffered saline (PBS) to 4 parts 30 % hydrogen peroxide to 1 part distilled water, with a final addition of one drop of 25 % ammonia per 10 ml of combined volume (Hemmi and Grunert, 1999; Coimbra et al., 2015a). Bleaching was carried out at room temperature and in the dark until the RPE had turned light amber to pale hazel in color, which usually took eight or more hours and could be carried out overnight. This slow bleaching method carried out at approximately neutral pH preserves the Nissl substance for the later staining of the retinal ganglion cells. Thereafter, the retina was washed in 0.1 M SPB three times, one minute each, to remove any residual bleach, then gently cleaned with soft, fine brushes, transferred onto a standard microscope slide, and mounted in a solution of 80 % glycerol and 20 % SPB with 0.1 % sodium azide added (Coimbra et al., 2015b). Two small strips of filter paper placed next to the retina served as spacers between wholemount and the coverslip. Finally, the edges were sealed with nail varnish to prevent evaporation. It generally took several days to clear the tissue in the glycerol (Coimbra et al., 2015a; Coimbra et al., 2015b).

Subsequent to the photoreceptor analysis, the retinal wholemounts were remounted and prepared for RGC analysis. The varnish seal around the coverslip was cut away and the retina carefully transferred to and washed in 0.1 M SPB three times, one minute each. Then, the retina was flipped over such that the RGC layer faced upwards, transferred to a Superfrost^®^ Plus adhesive microscope slide, and gently flattened without distending it. The slide was then placed overnight in a glass staining dish containing a few milliliters of 40 % formaldehyde solution to create an atmosphere saturated with formaldehyde vapor (Stone, 1981; Coimbra et al., 2006).

This procedure ensured the retina was well fixed and adhered to the slide. Subsequently, the retinal wholemount was stained for Nissl substances according to the protocol described in (Coimbra et al., 2006). Finally, the retinae were mounted in Eukitt^®^ mounting medium and were allowed to dry for at least 24 hours before analysis. This entire process can lead to shrinkage and distortions, especially when the retinal wholemounts (partially) detach from the microscope slide during the staining process. The retinae analyzed in this study did not detach during processing and shrinkage was very limited and corrected for in the following analysis.

### Stereological analysis and data processing

The retinal wholemounts were analyzed using a Zeiss AxioPlan II fluorescence microscope with a motorized specimen stage and a stereology system running Stereo Investigator 7 software from MBF bioscience. We used the optical fractionator method (West et al., 1991) with modifications for retinal wholemounts (Coimbra et al., 2009; Coimbra et al., 2012). The first stereological analysis was carried out with the glycerol-mounted, unstained wholemounts and focused on assessing the distribution of different retinal cone types within the photoreceptor layer. The receptor cells were visualized using a 40x magnification objective with differential interference contrast and focusing at the level of the cone inner segments. Individual rods could not be distinguished reliably throughout the retina due to their small size, the limited magnification available and tissue quality. However, all cone types could be readily distinguished by the size and shape of their inner segments. To obtain representative, systematic estimates of the density of each cone type and the total population of cone photoreceptors, we traced the coordinates of the entire retina’s outline and an additional outline that included the foveal and triple cone area. Both of these special retinal regions could be recognized at low magnification and we outlined them with a margin of 200 - 300 μm. To achieve sufficient areal coverage and representative sampling, we had the program assign 200 counting sites of 50 x 50 μm to the entire retina, which resulted in a grid width of 250 - 300 μm between loci, depending on the size of the retina. The foveal area was assigned a higher resolution grid with 100 - 150 μm between sites and counting frames of 25 × 25 μm to compensate for the marked changes in receptor density over small distances.

We repeated the same procedure for the RGC population within the retinal ganglion cell layer, using bright field illumination. We counted all cells with relatively large, slightly polygonal, positively Nissl-stained somata as RGCs, and only excluded cells with somata that were either distinctly small and round (amacrine cells) or cigar-shaped (glia). This approach has been found to provide an accurate representation of the ganglion cell population within the ganglion cell layer. However, despite the fact that in specialized regions, the high densities of neurons makes differentiating amacrine cells and ganglion cells difficult, the proportions of amacrine cells in these regions were found to be low and inclusion of amacrine cells in the counts does not affect the general topographic pattern (Collin and Pettigrew, 1988a).

The Stereo Investigator software automatically produces a summary of the collected counting data, including area sampling fraction, total population estimates, and coefficients of error (CE). We chose Scheaffer's CE to assess the accuracy of the obtained estimates, as it most closely approximates the true CE in a stereological sampling context (Glaser and Wilson, 1998). In biological contexts, sampling schemes that result in a CE < 0.1 are deemed sufficiently representative and accurate (Fileta et al., 2008). For further analysis, the raw data were exported from Stereo Investigator to JMP^®^ v.11.1.1 (SAS Institute Inc.). Some counting sites (5-10 per retina) were randomly allocated to locally compromised parts of the wholemounts. This mostly affected single cone counts, while the larger and characteristically shaped double cones usually were still discernible. In such cases, the cone ratio of neighboring sites and the double cone counts from the ambiguous site were used to correct the corresponding single cone numbers. If neither cone type was discernible, the entire site was omitted from counting (1-3 per retina). To correct the RGC density values for shrinkage due to Nissl staining, we divided the retinal wholemounts digitally into distinct regions, measured their area before and after staining in ImageJ v.1.48t, and calculated the individual shrinkage factor for each region. The raw RGC densities from each counting site were then multiplied with their respective region's shrinkage factor. The corrected data were converted into topographic maps by plotting the densities of the individual cone types and RGCs against the X- and Y-coordinates of the counting sites, using JMP's contour plot tool. These contour plots were then exported to Microsoft^®^ PowerPoint^®^ (v.14.2.3 for macs) and further edited, e.g. re-matched with their respective retinal wholemount's original outline. The approximate angular position of the fovea was estimated from the finished topographic maps in ImageJ v.1.48t, based on its relative horizontal and vertical distance from the center of the retina, and assuming the horizontal and vertical diameter of the retina correspond to 180°.

### Calculating spatial resolving power

We based our estimate of *T. delaisi*'s anatomical spatial resolving power (SRP), expressed as Nyquist frequency (*fN*) in cycles per degree (*cpd*), on the eyes' posterior nodal distance (*PND*, in mm) and foveal photoreceptor cell density (*D*, in mm^−2^), according to the following equation, modified from Williams (1988):

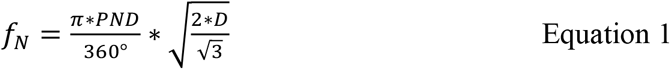

Due to the lateral displacement of RGCs in foveae (Wassle et al., 1990), we assumed the photoreceptor cell density to be the most reliable estimator for the anatomical spatial resolving power. Our reasoning will be explained in more detail in the results section. To reflect the potential of this species, we used the highest individual density value. A previous study on paired cones suggested that each cone member could produce an individual signal in the context of color vision (Pignatelli et al., 2010). If this is confirmed, individual signals might also be used for greater resolution. We therefore calculated both a conservative estimate for the anatomical SRP, assuming each twin cone to signal as a unit, and a maximal estimate, assuming each twin cone member signals individually, to show the highest possible SRP for this species. The linear density is based on a hexagonal cell arrangement in the fovea, which accounts for the 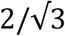 factor in equation 1.

## Results

### General eye features

*T. delaisi* possesses relatively large and conspicuous eyes for its small body size (Fig. 1A). While standard length averaged 42.7 ± 2.8 mm (mean ± SD, *N* = 10), the eyes had diameters of 3.48 ± 0.15 mm antero-posteriorly and 3.00 ± 0.15 mm dorso-ventrally, corresponding to 8.2 ± 0.5 % and 7.0 ± 0.4 % of the standard length. Eyecup and retina are therefore hemi-elliptical with the dorso-ventral axis measuring 86.3 ± 3.8 % of the length of the antero-posterior axis. This causes the distance between the spherical lens and different regions of the retina to vary and has implications for image focusing that are discussed later. The pupil of *T. delaisi* also features an elliptical shape (Fig. 1B), with the antero-posterior axis measuring 1.47 ± 0.09 mm and the dorso-ventral axis measuring 1.24 ± 0.08 mm. This creates an antero-ventral aphakic gap when the lens is in its relaxed position, *i.e*. its focus set to infinity. The lens diameter measured 1.40 ± 0.08 mm (N = 4) in the eyes dissected for the retinal wholemounts, closely matching overall average pupil size. A well-developed *stratum argenteum* lines the back of the eye as part of the choroid, between the retinal pigment epithelium and the sclera. Apart from containing the defining silvery, and highly reflective stacks of guanine crystals, the *stratum argenteum* is also associated with yellow chromatophores that give it a golden sheen. The *argenteum* was found to be continuous with a layer of iridophores in the iris that contained similar stacks of crystals but lacked the yellow pigment granules.

**Fig. 1:**
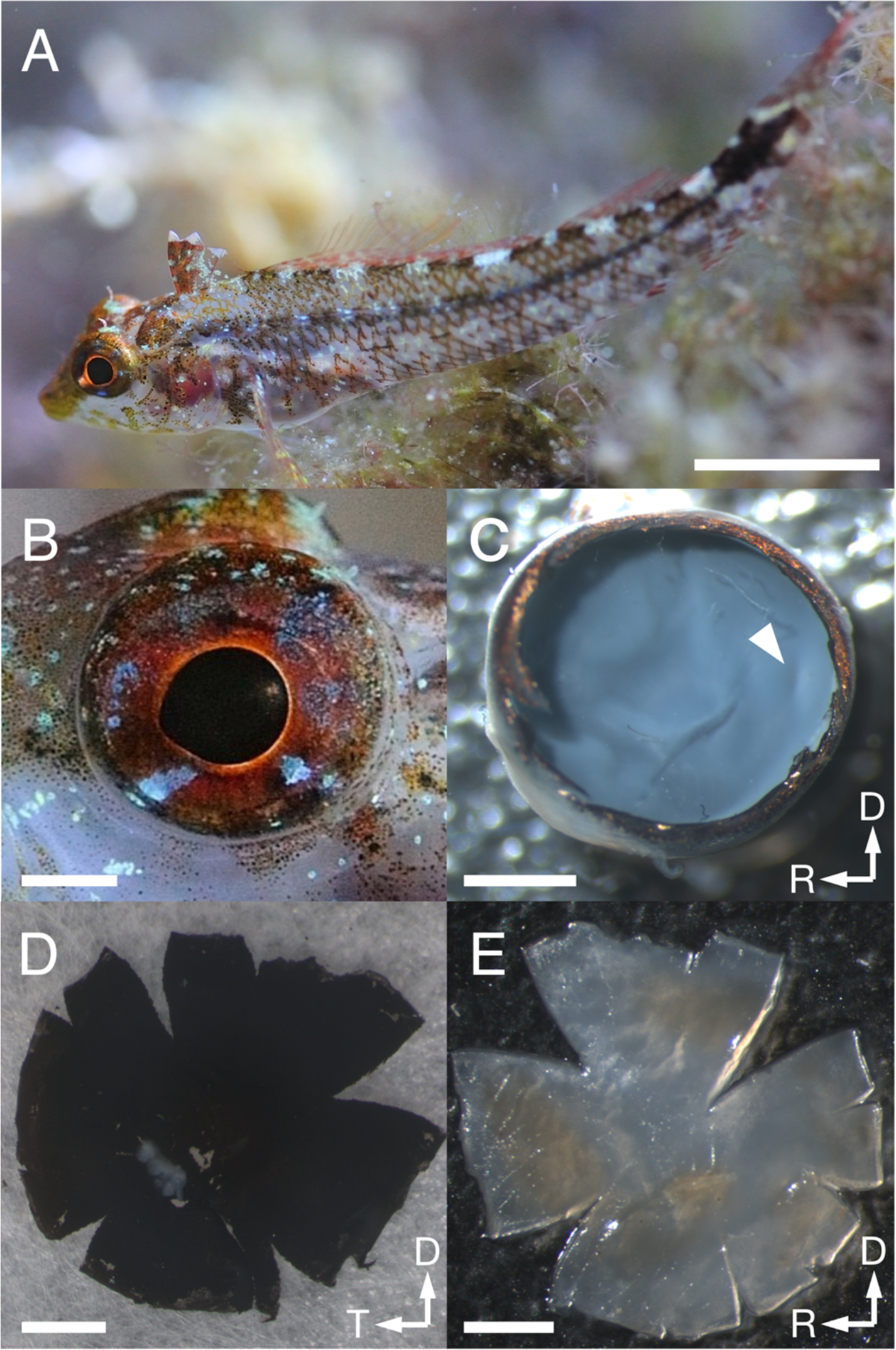
*T. delaisi* habitus and eye preparation. (A) *T. delaisi* in its natural environment. (B) Close-up of the eye showing the slightly elliptical dimensions of eyeball and pupil. (C) After removing cornea, iris, and lens, the remaining eyecup was fixed until the retina took on the translucent appearance seen here. The arrowhead marks the macroscopically visible, slightly elongated foveal pit situated in the temporal retina. (D) Back of the isolated retina with retinal pigment epithelium still attached. (E) Retina after bleaching and flattening, seen from the vitread side, ready to be mounted on a slide. Scale bars equal 1 cm in A, and 1 mm in B-E; arrows indicate dorsal, D, rostral, R, and temporal, T, directions.

*T. delaisi*'s retina covers an almost hemispherical visual field. The horizontal viewing angle approximately spans 160°, as estimated from horizontal eye sections. In addition, *T. delaisi* is able to rotate its eyes by 60-70° in the horizontal plane. Consequently, the triplefin can observe its entire surrounding by eye movement alone and has areas of binocular overlap in frontal and caudal direction (Fig. 2). Due to our limited data on the subject, the extent of *T. delaisi*'s visual field and eye movements shown here may even be an underestimate and has to be validated in future studies.

**Fig. 2:**
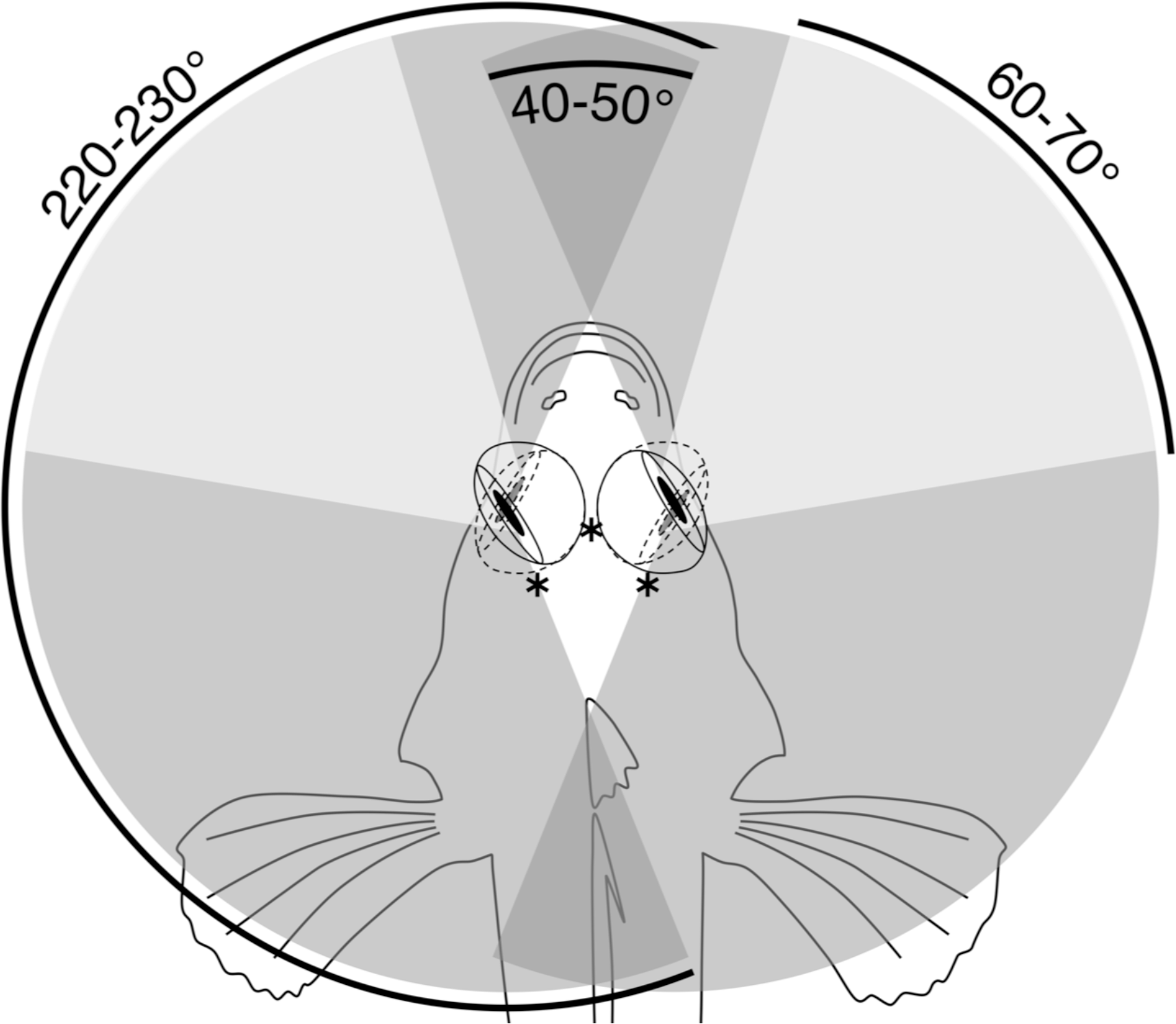
Horizontal visual field in *T. delaisi*. Schematic top view of *T. delaisi*'s head showing the extreme frontal and temporal orientation of its eyes (solid and dashed lines) and the associated foveal positions (*). The shaded areas indicate the resulting maximum total (medium grey), binocular (dark grey), and foveate (light grey) horizontal fields of view based on the extent of eye movement.

### Photoreceptors

We found four morphologically distinguishable photoreceptor types: rods, single cones, paired cones, or more specifically twin cones due to their morphologically equal members, and triangularly arranged triple cones with morphologically identical members. They occurred in three distinct mosaic patterns (Fig. 3). Rods outnumbered cones in the periphery (Fig. 3A) but occurred only outside the foveal area. They were not explicitly counted because their small diameter did not allow for reliable identification within our sampling framework. Single and twin cone densities were generally higher in the dorsal and temporal retina than in the ventral and nasal retina. The continuous increase in density from the periphery to the dorso-temporal retina culminated in a pronounced *area centralis* with a distinct fovea (Figs. 4 and 5; detailed data in Table 1). Single cone density averaged 10,200 cells per mm^2^ and peaked at 30,800 cells per mm^2^ (typical distribution shown in Fig. 4A). Twin cones averaged 23,300 cells per mm^2^ and peaked at 104,400 cells per mm^2^ (typical distribution shown in Fig. 4B). With a total retinal surface area of 16.09 ± 0.18 mm^2^ (Table 1), these densities resulted in average total population estimates of 165,700 single cones and 369,700 twin cones for the entire retina. Triple cones were only found in a limited retinal region between the optic disc and the fovea (Fig. 4C) at an average density of 16,700 ± 2,500 cells per mm^2^ (Table 1). At their peak density of 30,400 ± 5,700 cells per mm^2^, they outnumbered both single and twin cones in the same area. Due to their restricted distribution, the total number of triple cones was estimated at only 5,350 ± 1,100 (Table 1). The sum of all cone types showed a continuous distribution and consistent increase in overall density towards the fovea (Fig. 4D). More stereological data and Scheaffer's coefficient of error (CE) for each cone type are listed in Table 1. The relatively high CE for the triple cone estimates can be attributed to their restricted distribution and correspondingly fewer counting squares.

**Fig. 3:**
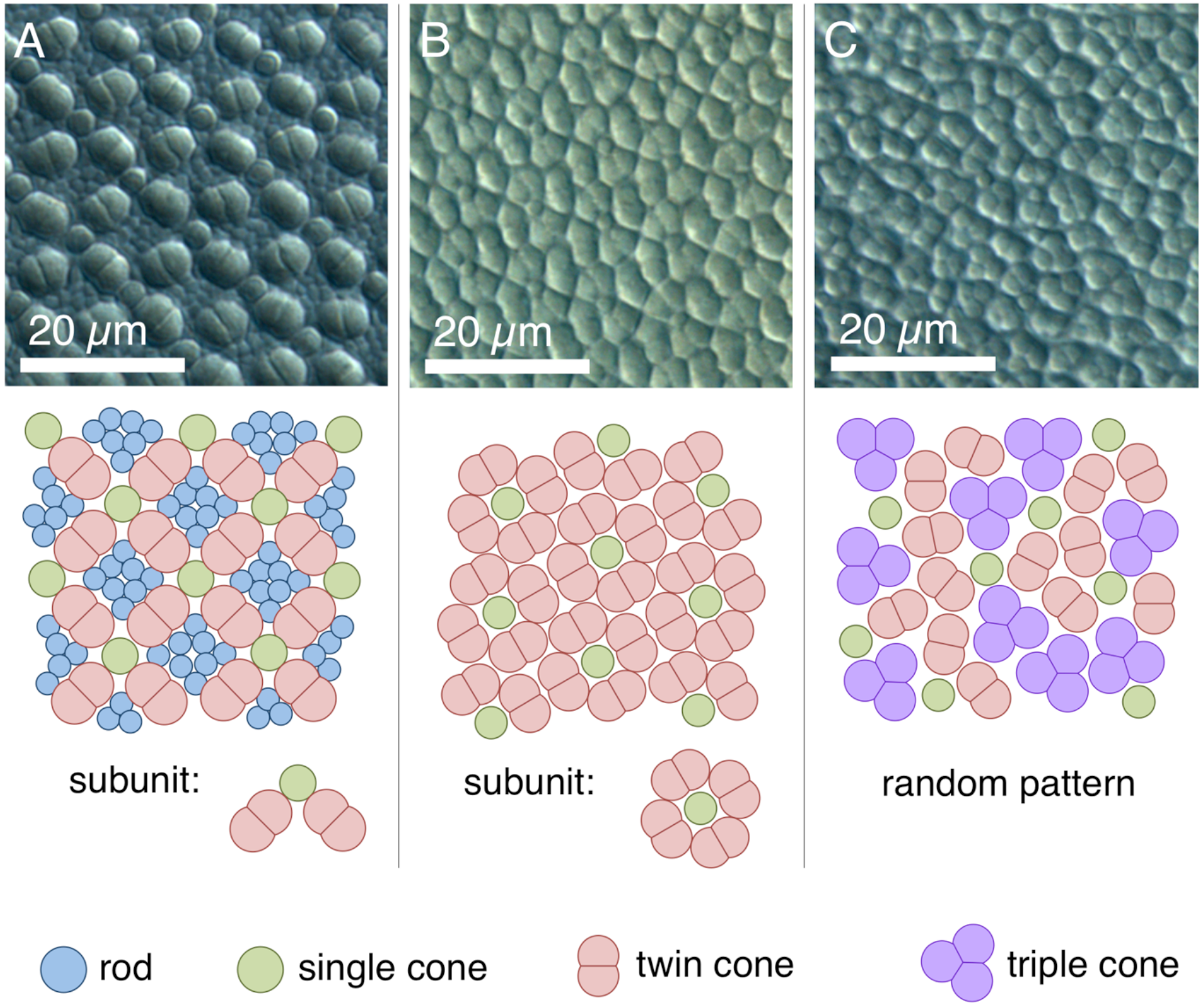
Cone types and mosaic patterns in the retina of *T. delaisi*. (A) Square mosaic pattern with a 2:1 ratio of twin to single cones, as found across most of the retina. Rods fill in the gaps between cones. (B) Dense, "lucky-clover" pattern with 4:1 ratio of twin cones to single cones and without rods, as found around the fovea. The fovea itself features a modified version of this pattern (Fig. 5D). (C) A distinct area between optic disc and fovea features triple cones and a random arrangement of all cone types.

**Fig. 4:**
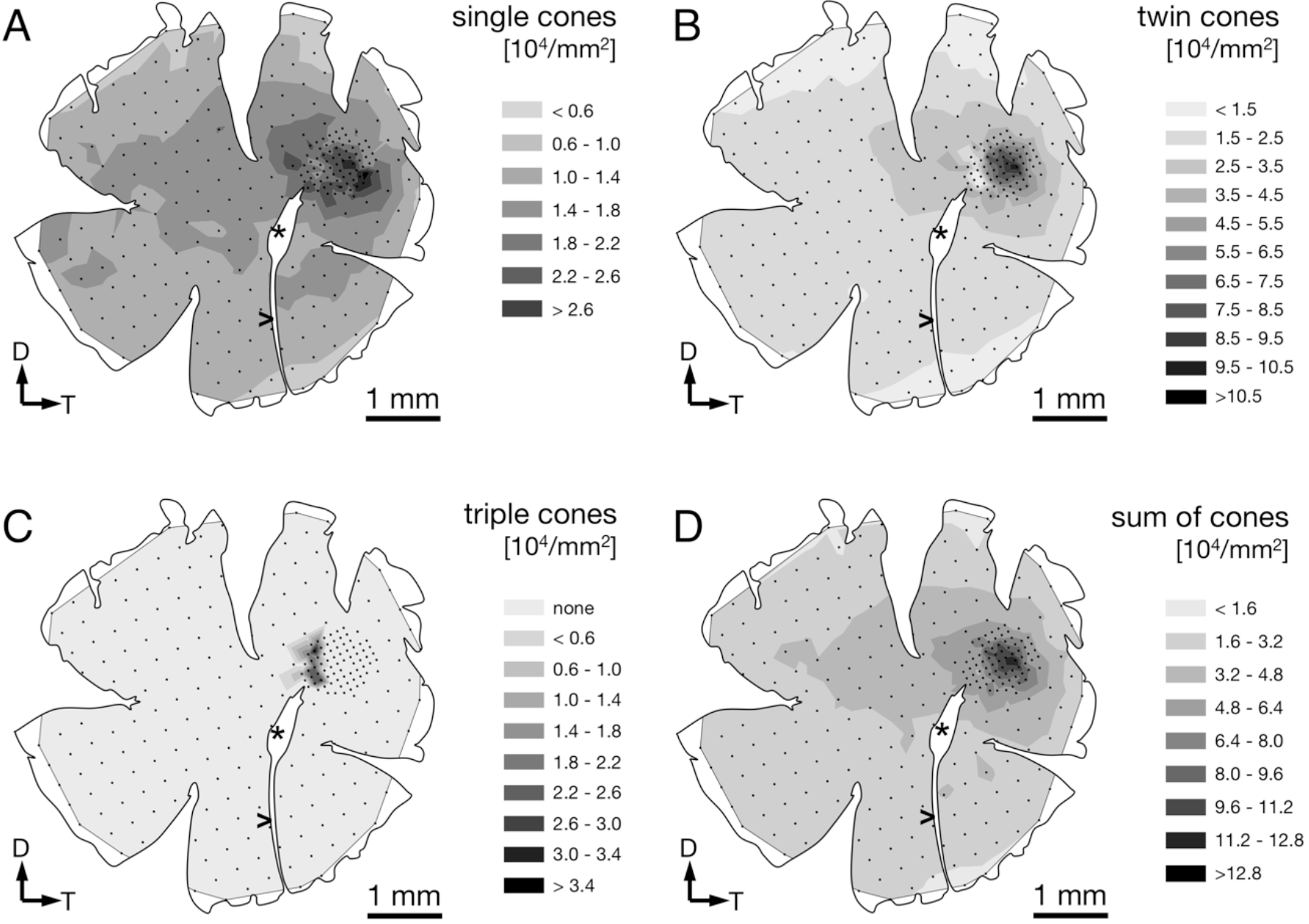
Retinal topography of different cone types in a representative individual. The maps show the left retina of one *T. delaisi* individual. The central hole and the gap extending nasoventrally are from the optic nerve head (*) and falciform process (>). Black dots in the map correspond to counting sites. Arrows indicate dorsal, D, and temporal, T, direction.

**Fig. 5:**
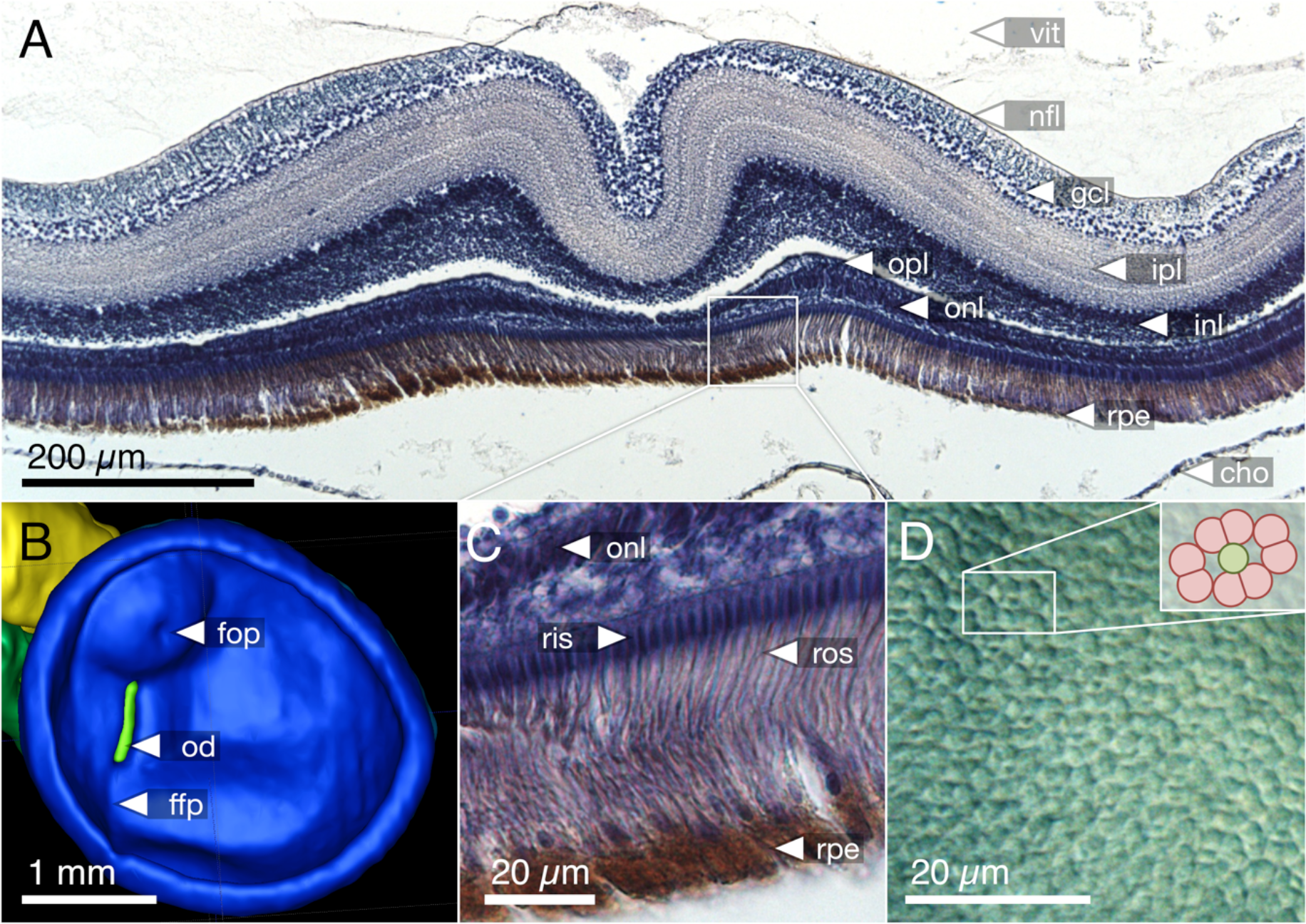
Foveal features in *T. delaisi*. (A) Coronal section of paraffin-embedded retina showing a steep foveal pit and locally thickened retina. The gaps between the outer plexiform and inner nuclear layers, as well as the retinal pigment epithelium and choroid, are artifacts of the tissue processing. (B) Virtual segmentation of an *in situ*, right-eye retina from magnetic resonance imaging data. The foveal area forms an almost hemispherical bulge in the retina, confirming the histological findings. The pronounced semi-ellipsoid shape of the retina matches that of the external eye. (C) Magnified view of the inset in (A). Note the densely packed receptor inner and outer segments with a diameter of about 2 μm each. (D) Top view of the cone mosaic pattern in the fovea. The inset demonstrates that it is a skewed version of the 4:1 pattern shown in Fig. 3B. cho = choroid; ffp = falciform process; fop = foveal pit; gcl = ganglion cell layer; inl = inner nuclear layer; ipl = inner plexiform layer; nfl = nerve fiber layer; od = optic disc; onl = outer nuclear layer; opl = oter plexiform layer; ris = receptor inner segments; ros = receptor outer segments; rpe = retinal pigment epithelium; vit = vitreous body.

**Tab. 1:** Summary of stereological parameters and data, presented as means ± SD based on *N* = 4 individuals. "N sites" includes only sites with actual counts for the respective cell type (mean ± SD, rounded to integer sites). CE, coefficient of error.

| cell type | sampling region | area (mm<sup>2</sup>) | N sites | area sampling fraction | Scheaffer's CE | mean density (mm<sup>-2</sup>) | max. density (mm<sup>-2</sup>) | total population |
| --- | --- | --- | --- | --- | --- | --- | --- | --- |
| single cones | general retina | 15.14<br>± 2.31 | 167<br>± 21 | 0.0276<br>± 0.0027 | 0.020<br>± 0.006 | 10,200<br>± 600 | 30,800<br>± 7,300 | 165,700<br>± 15,000 |
| | foveal area | 0.95<br>± 0.18 | 33<br>± 11 | 0.0222<br>± 0.0082 | 0.082<br>± 0.042 | | | |
| twin cones | general retina | 15.14<br>± 2.31 | 175<br>± 12 | 0.0292<br>± 0.0030 | 0.020<br>± 0.002 | 23,300<br>± 2,700 | 104,400<br>± 25,400 | 369,700<br>± 30,700 |
| | foveal area | 0.95<br>± 0.18 | 46<br>± 9 | 0.0315<br>± 0.0111 | 0.054<br>± 0.014 | | | |
| triple cones | triple cone area | 0.32<br>± 0.03 | 14<br>± 2 | 0.0265<br>± 0.0026 | 0.225<br>± 0.130 | 16,700<br>± 2,500 | 30,400<br>± 5,700 | 5,350<br>± 1,100 |
| ganglion cells | general retina | 15.19<br>± 1.81 | 180<br>± 19 | 0.0299<br>± 0.0038 | 0.025<br>± 0.002 | 32,400<br>± 2,000 | 81,000<br>± 10,300 | 516,800<br>± 38,600 |
| | foveal area | 0.90<br>± 0.35 | 45<br>± 14 | 0.0322<br>± 0.0058 | 0.021<br>± 0.008 | | | |

In terms of areal coverage, the retina of *T. delaisi* was found to be cone-dominated throughout, with an estimated 75 - 80 % of its area occupied by cones (Fig. 3). Most regions of the retina featured a cone mosaic pattern with a ratio of two twin cones to one single cone. The twin cones were arranged in a regular grid-like pattern with the single cones in the corner positions of the squares (as defined by Lyall (1957); see also Fig. 3A). We refer to this as the 2:1 square mosaic. This cone pattern was interspersed with randomly arranged rods in the spaces between the cones. Near the fovea, the cones became slenderer and were arranged more densely, while rods gradually decreased in number. In the area immediately surrounding the fovea, rods were missing entirely. The prevailing pattern changed such that the single cones were now in the central position of the twin cone squares and none of the surrounding twin cones was shared with other single cones (Fig. 3B). This resulted in a 4:1 ratio of twin to single cones and is thus referred to as the 4:1 square mosaic. The fovea featured a skewed version of the 4:1 mosaic, in which twin cones formed a diamond rather than a square pattern (described below, Fig. 5D). The triple cone area featured all cone types in an irregular arrangement, and no detectable rods (Fig. 3C).

### Fovea

The fovea in *T. delaisi* was apparent even macroscopically during the dissection of the eyes and preparation of the retinal wholemounts and appears as a thickened area with a central dip in the temporal region of the retina. The three-dimensional shape of the fovea is distorted by flattening in retinal wholemounts. The histological sections (e.g. Fig. 5A) revealed the thickened retinal layers in the foveal area, and the central dip. Possibly due to differential shrinkage, however, the retina separated from the choroid and sclera, probably compromising the foveal shape in the process. We consider the virtual segmentation images produced from magnetic resonance imaging (MRI) data the most reliable impression of the foveal shape, as these were obtained from whole eyes *in situ*, and they revealed the foveal area to form an almost hemispherical convexity (Fig. 5B). The position of the fovea, as estimated from the retinal wholemounts and in relation to the center of the retina, was approximately 30° ± 5° (*N* = 4) dorsal and 50° ± 9° (*N* = 4) temporal. This foveal position, when projected onto the visual field, means that *T. delaisi*'s high-acuity vision, and thus visual focus, is directed about 40° sideways and 30° downward in relation to its body axis when the eye is in its resting position. High-amplitude eye movements, however, allow the fovea to be pointed towards a large part of the frontal visual field and to image any object therein with high resolution (Fig. 2). Nevertheless, eye movement seems to create only approximately 40° of binocular overlap, which may not include the foveal visual field. The apparent dip in the thickened foveal area was confirmed to be the vitread opening of a deep foveal pit with steep, convexiclivate walls (Fig. 5A). Some section series suggested that the foveal pit was actually formed like a trough between the fovea and the optic nerve head, possibly allowing the axons of foveal ganglion cells to run to the optic nerve head along the shortest possible path. This could not be ascertained in all sections. The highest individual receptor cell density that we found in any fovea was 160,000 cones per mm^2^, comprised of 132,000 twin cones per mm^2^ and 28,000 single cones per mm^2^. This density was achieved through a gradual, marked reduction in cone cell width from the periphery towards the fovea and the omission of rods in the foveal area (Figs. 3 and 5C–D). The above density theoretically leaves an average area of 2.5 × 2.5 μm^2^ for each cone to occupy in the fovea, which was matched by the 2 - 3 μm diameters of the cone inner segments (Fig. 5C-D).

### Retinal ganglion cells

Demounting, flipping, Nissl-staining, and remounting the retinas were successfully performed without damaging the wholemounts, which allowed us to determine the retinal ganglion cell (RGC) distribution in the same retinae for which we had already obtained the receptor cell densities (Fig. 6A). The stained RGC nuclei could be distinguished from other cell nuclei in the same retinal layer by their characteristic, slightly polygonal shape (Fig. 6B).

**Fig. 6:**
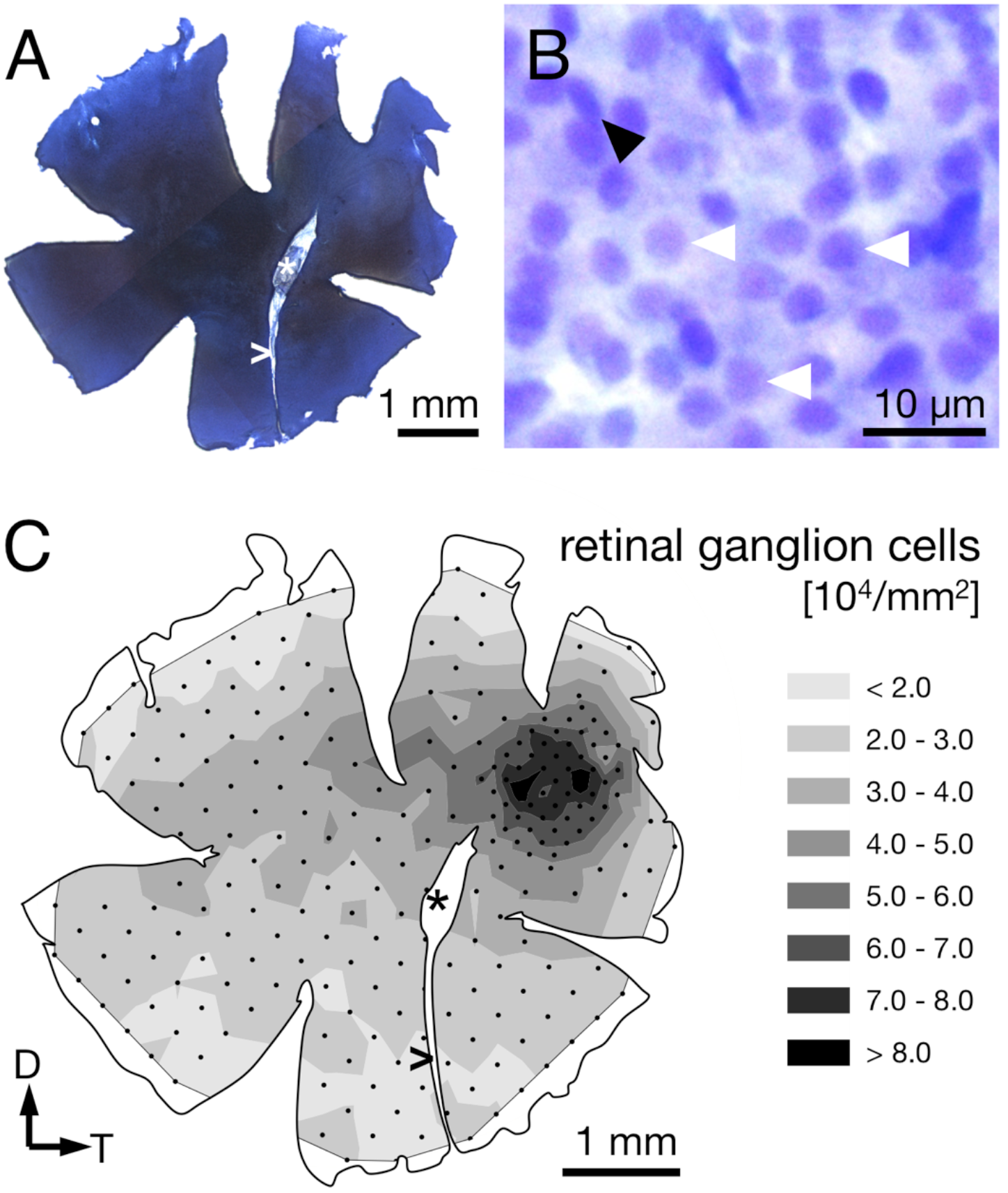
Ganglion cell counts from Nissl-stained retinae. (A) Sample retinal wholemount after Nissl-staining. Note that this is the same left retina for which receptor cell densities were presented in Fig. 4. Apart from limited shrinkage along the edges, its size and outline are well preserved. (B) The retinal ganglion cell layer at 630x magnification. Ganglion cells feature relatively large, slightly polygonal nuclei (white arrowheads). Smaller or elliptical nuclei (black arrowhead) were attributed to different cell types and not counted. (C) The density distribution of retinal ganglion cells matches the overall pattern found for the photoreceptors. Note that the highest densities are found in the perifovea, with a drop in density in the center of the fovea. Black dots correspond to counting sites. Arrows indicate dorsal, D, and temporal, T, direction, * marks the optic nerve head, and > the falciform process.

The RGC showed an overall mean density of 32,000 ± 2,000 cells per mm^2^ (Table 1). Their distribution followed that of the photoreceptor cells in that it was generally higher in the dorsal than the ventral part of the retina and steadily increased towards the fovea (compare Figs. 4D and 6C). Maximal RGC densities occurred in a ring surrounding the fovea and reached 81,000 ± 10,300 RGC per mm^2^ (Fig. 6C). The center of the fovea featured high numbers of RGC as well, but their density decreased slightly compared to the perifovea, which is the result of the thickened retinal layers in this area and the shape of the foveal pit. Stereological parameters and coefficients of error can be found in Table 1.

### Anatomical spatial resolving power

The lateral displacement of RGCs in the foveal area makes it impossible to infer the actual connectivity directly from the apparent RGC density. The center of the primate fovea features few, if any, RGCs, and yet each foveal photoreceptor is connected to 3 - 4 displaced RGCs in the perifovea (Wassle et al., 1990). In *T. delaisi*, we found the peak RGC densities located in the perifovea and decreasing towards the center of fovea, which suggests a similar lateral displacement, albeit to a lesser extent. Additionally, the fovea is generally accepted to be a retinal specialization for high-resolution vision, and therefore it is reasonable to assume the circuitry in the fovea is adapted to allow maximal resolution. For these reasons, we assumed a connectivity ratio of at least 1:1 between foveal photoreceptors and RGCs in *T. delaisi*, and used its photoreceptor cell densities directly to calculate the anatomical spatial resolving power (SRP). In the more conservative approach we used the highest individual combined density of single and twin cones, counting the latter as one receptor, which yielded a density of 160,000 per mm^2^. For the maximal anatomical SRP estimate, the assumption that each twin cone member provides direct input (Pignatelli et al., 2010), lead to a maximal receptor cell density of 292,000 per mm^2^. We estimated the posterior nodal distance (PND) at 1.79 mm, based on *T. delaisi*'s lens radius of 0.70 ± 0.04 mm (*N* = 4) multiplied by 2.55, the average Matthiessen's ratio for teleost eyes (Shand et al., 1999). The resulting anatomical SRP was 6.7 cpd under the conservative assumptions and 9.0 cpd under the maximal assumptions. These values translate to a minimal separable angle (MSA) of 0.15° and 0.11°, respectively. Such MSAs would allow *T. delaisi* to resolve a 1 mm sized object at a distance of 38 - 52 cm (see Fig. 7 for an ecological example).

**Fig. 7:**
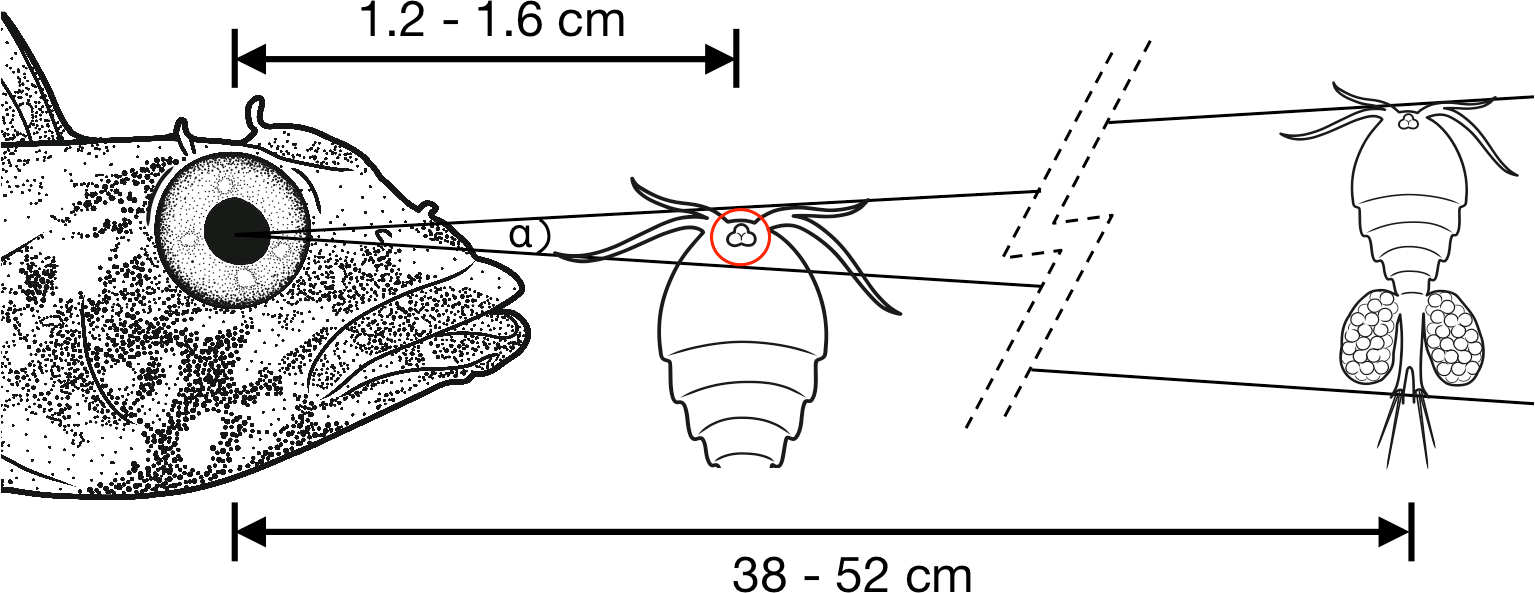
Illustration of the minimal separable angle (MSA; α) in *T. delaisi*. With its estimated MSA of α = 0.11° - 0.15°, *T. delaisi* could resolve a 1 mm long copepod, a typical prey item, at a distance of 38 - 52 cm. The eye of the copepod could play a role in its detection or identification as prey (Harant and Michiels, 2017). Assuming a diameter of 30 μm in this example, it might be resolved at 1.2 - 1.6 cm. The required ability to focus at such short distances is facilitated by several features of *T. delaisi*'s eyes. Sizes, distances, and angles are not to scale.

## Discussion

Within the large number of studies on fish retinal topography (Collin, 2008) and visual acuity (Caves et al., 2017), there is only little and incomplete information on small, crypto-benthic fishes. By utilizing the retinal wholemount technique, complemented by additional histological and MRI data, this study provides a comprehensive insight into the visual morphology of the micro-predatory triplefin blenny *Tripterygion delaisi*.

## General eye anatomy

Like most small, benthic fishes, *T. delaisi* possesses prominent eyes in a pronounced dorsal position, protruding from the head. In combination with their behavior of rearing up their anterior body half in their resting position (Wirtz, 1978), *T. delaisi* will have a slightly raised viewing angle of their surroundings, which might help to avoid their view getting blocked by algae or other substratal structures. The relatively large size of *T. delaisi*'s eyes is common among small fishes. The correlation between body size (measured as total length) and eye diameter in 61 teleosts reported by Caves et al. (2017) suggests that *T. delaisi*'s eyes are actually slightly smaller than expected for a fish of its size. However, total length might not be an accurate predictor in species with long and slender bodies. An important factor limiting eye size is head width. Since *T. delaisi*'s eyes are almost touching medially, they have maximized eye size within the constraints of their head size and shape, underlining the importance of vision to them.

Fish eyes that require a wide accommodative range can adapt in different ways. In most cases the eyes simply evolve to be longer along the axis of the accommodative lens movement, whose largest component is usually along the pupillary plane (Fernald, 1990). Accordingly, the reason for the large relative distance between lens and retina in sea horse eyes (Mosk, 2004) could be the animals' need to bring micro-prey directly before their snouts into focus, since they are visual planktivores (Lee and O'Brien, 2011). The sandlance Limnichthyes fasciatus also requires precise and wide-range accommodation, as it strikes after tiny planktonic organisms that drift by at different distances (Pettigrew et al., 2000). In contrast to sea horses, it evolved an optical system unique among teleosts, with an adjustable corneal lenticle and a flattened lens, which, in combination, have an exceptionally high accommodative power and range (Collin and Collin, 1988a; Collin and Collin, 1988b; Pettigrew and Collin, 1995). *T. delaisi* widens its accommodative range with an elliptical eyeball. The longer naso-temporal axis improves their ability to focus close objects in the anterior visual field. In order to maintain their own camouflage, triplefins minimize body movement, and hence only snatch at prey in their immediate surroundings. The ability to form a sharp image at very short distances is crucial for both detecting small and well-camouflaged prey against a complex substrate, in order to reduce striking-distance without relinquishing precision. A consequence of the lens moving parallel to the pupil is the coupling of accommodative states of different retinal areas. When the temporal retina is focused on the foreground in the anterior field of view, the nasal part must be focused on the background of the posterior field of view, and *vice versa*, while the central part of the retina is restricted to a constant, intermediate accommodative state (Fernald, 1990). This could be an advantage, as it enables *T. delaisi* to simultaneously have a focused image of both prey directly in front, and potential predators behind, while they are still at a distance.

The elliptical shape of the pupil results in an aphakic space, where light can enter the eye without passing through and being focused by the lens. Several non-exclusive functions have been suggested to explain this ocular specialization. Rostral aphakic spaces, like the one in *T. delaisi*, are common in diurnal fishes (Schmitz and Wainwright, 2011) and have been suggested to extend the anterior visual field and to allow greater binocular overlap (Walls, 1942). The potential binocular overlap in *T. delaisi*, however, is small, and might even be irrelevant in this species, as discussed later. Another function, especially for circumlental aphakic gaps, could be increasing retinal illumination (Munk, 1980; Warrant and Locket, 2004). Although *T. delaisi* inhabits micro-environments with reduced or fluctuating light levels within the photic zone (Wirtz, 1978), these are still within the range of photopic vision, making an increase of general retinal illumination unlikely to be advantageous during the day. The most parsimonious functional assumption for the aphakic gap is therefore the provision of the required room for the accommodative movement of the lens, which protrudes through the pupil in fishes (Sivak, 1978; Nicol and Somiya, 1989). Accordingly, the location of the aphakic gap depends on the accommodative state of the eye and its orientation correlates with the direction of lens movement, which commonly coincides with the visual axis of highest acuity (Nicol and Somiya, 1989). This applies to *T. delaisi* as well, allowing it to bring objects from a wide range of distances into focus where it can see them at its highest spatial resolution, which is an essential adaptation for a micro-predator that needs to detect small, hidden prey in its vicinity.

### Eye mobility and visual field

Another indicator of the importance of vision for *T. delaisi* is the ability of moving its eyes around all axes of rotation and largely independent of each other. The degree of independent eye movement varies considerably in other fish species (Fritsches and Marshall, 2002). In this respect *T. delaisi* probably lies between syngnathids, with somewhat independent eyes, and the sandlance, with completely independent eye movements. The lateral position and high mobility of *T. delaisi*’s eyes result in a horizontal visual field of about 220° - 230° for each (Add. Fig. 1). Triplefins constantly use this eye mobility to monitor their entire surroundings, while keeping the body still. This behavior complements the triplefins' cryptic body coloration and maximizes their camouflage, allowing them to see, while remaining unseen. Furthermore, elaborate and frequent eye movements can be indicative of a localized zone of acute vision, like an area centralis or fovea, in the retina of any species (Bartels, 1931; Kahmann, 1936), an assumption that is met by *T. delaisi*. The dorso-temporal location of *T. delaisi*'s fovea puts the zone of highest acuity in the naso-ventral visual field, directed towards the substrate in front of the fish. This can be seen as an adaptation to the triplefin's targeting benthic invertebrates, and is generally expected for predators of substratal, small prey (Nicol and Somiya, 1989). Based on the estimated extent of eye movements, 60° - 70° horizontally, foveate vision can cover a total of about 120-140° in the anterior visual field (Add. Fig. 1). The maximal binocular overlap, however, appears to be only about 40° - 50°, which is not enough to include foveate stereoscopic vision. *T. delaisi*'s independent eye movement and its habit to focus on prey with just one eye before striking sideways, suggest that binocular foveate vision may be irrelevant to them and further imply a monocular distance estimating mechanism. Similar findings were reported for pipefish, including a possible functional link between their steep foveal pit and monocular depth perception (Collin and Collin, 1999).

### Photoreceptor types and mosaic

*T. delaisi*'s retina was found to be cone-dominated, containing single, twin, and triple cones, while the number of rods declined from the periphery towards the fovea. Such cone-dominated retinae are common for diurnal, coastal fish species (Bowmaker, 1990). The wavelengths of maximal absorption of *T. delaisi*'s different photoreceptor types are about 500 nm for rods, 470 nm for the single cones, and 516 / 530 nm for the two members of the twin cones (Bitton et al., 2017). Possessing two different visual pigments while being morphologically indistinguishable, *T. delaisi*'s paired cones fall into the category of non-identical twin cones (Nicol and Somiya, 1989). The maximal absorption values lie within the variation of visual pigments across species, but their range is narrower and shifted towards the green part of the visible spectrum, compared to the average sensitivities found in other species (Bowmaker, 1990; Losey et al., 2003). Individual double cones members have been shown to produce independent signals and thus enable trichromatic color vision in a species of triggerfish (Pignatelli et al., 2010). The same could be true for *T. delaisi* and allow enhanced discrimination of green and yellow colors, which are predominant in the triplefin's sublittoral environment of algae-and sponge-encrusted substrates. Assessing visual capabilities should not be reduced to the receptors' absorption maxima alone, however, but the broad shape of the visual pigments' absorption spectra, which extends *T. delaisi*'s color perception well into the red part of the visible spectrum (Bowmaker, 1990; Kalb et al., 2015; Bitton et al., 2017). The triple cone related findings of this study are discussed in a separate section below.

Most teleosts have duplex retinae that contain both rods and cones, responsible for scotopic and photopic vision, respectively (Nicol and Somiya, 1989). The decrease of rod density and cone size from the peripheral retina to the fovea in *T. delaisi* can be seen as a compromise between adaptation for sensitivity in the periphery and for acuity in the foveal area. The photoreceptors of most teleosts are arranged in regular patterns, whose many types vary with phylogeny, ontogeny, and location across the retina (Lyall, 1957; Engstrom, 1963; Wagner, 1972). The framework of this so-called mosaic is formed by the cones, while the rods, if present, are usually randomly arranged and inserted between the cones. The dominant cone mosaic found throughout most of *T. delaisi*'s retina is a square mosaic with each single cone being surrounded by four twin cones, which it shares with the neighboring single cones, thus resulting in a 2:1 ratio between twin and single cones (Fig. 3A; Fig. 8). The mosaic is unusual in that all single cones throughout the retina are in the corner position ("additional cones") and never in the central position, as defined by Lyall (1957) based on the orientation of the paired cones. This pattern variant has only been previously observed in *Nannostomus eques* and *Blennius vulgaris* (Wagner, 1972). To our knowledge, the lucky-clover-like pattern with each single cone surrounded by its own four twin cones (Fig. 3B) within the *area centralis* has not been described for any other species.

**Fig. 8:**
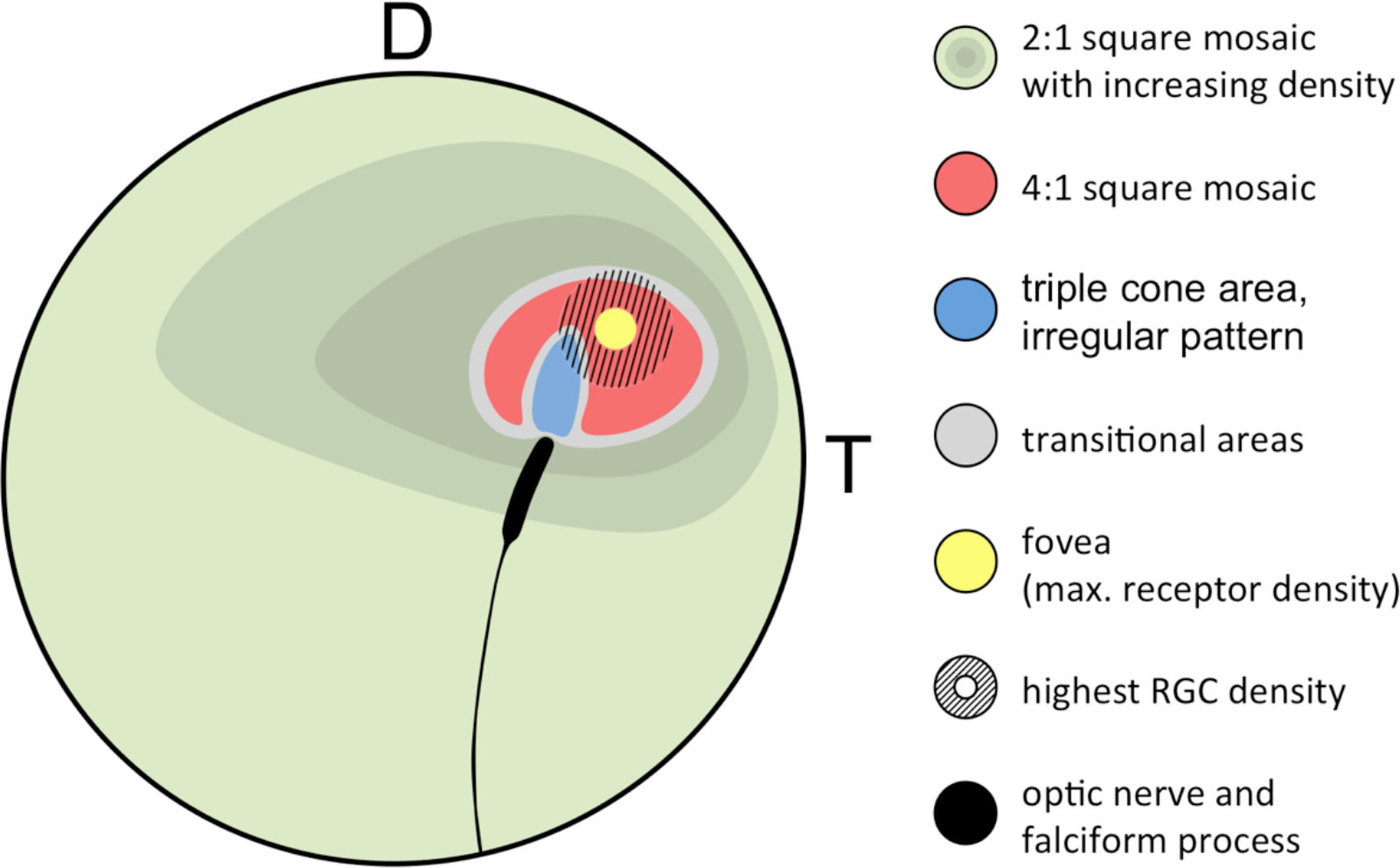
Generalized retinal map of *T. delaisi*. The illustration summarizes the relative position and covered area of cone density, mosaic patterns, and other features found in the retinae of four *T. delaisi*. D = dorsal, T = temporal, RGC = retinal ganglion cell.

The role of a regular cone mosaic is still uncertain and has never been tested experimentally, but several functions have been suggested based on correlations between pattern types and the ecology of multiple species (Wagner, 1990). Possibly the simplest explanation would be the maximization of receptor packing. Geometrically analyzing different possible patterns and especially paired cone shapes showed that in the most common mosaic patterns the cones achieve almost maximal areal coverage, which would increase photon catch and sensitivity at a given density (van der Meer, 1992). This seems to apply to the 4:1 square mosaic surrounding the fovea, as there are no gaps left between the cone inner segments. The 2:1 square mosaic in most of the retina has obvious, large gaps between the cones, but these are filled with rods, such that the areal coverage is also maximized. In the center of the fovea, however, maximal packing is not realized, as there seem to be actual gaps between the cone inner segments. The gaps might be due to the changed twin cone shape, *i.e*. the twin members are rounder and relatively further apart, combined with maintaining the same regular 4:1 arrangement. This could potentially indicate that the preservation of the mosaic is either a physiological constraint, or that mosaic regularity is more important than achieving maximum density in *T. delaisi*.

Another suggested function of cone mosaics is the improvement of motion detection through a dense and symmetrical arrangement of photoreceptors (Lyall, 1957; van der Meer, 1992). Square mosaics occur more often in species moving, or observing movement, in all three dimensions of their habitat, while row mosaics are more common in schooling fishes with predominantly horizontal swimming movements (Wagner, 1990). The consistent regularity of *T. delaisi*'s cone mosaic throughout most of its retina could grant it a sensitive and accurate sense of motion perception in all directions and across its entire visual field, which would benefit its predation of small but sometimes fast moving crustaceans in a complex micro-environment. Acute motion detection might also allow the small flicking movements of their first dorsal fin to be an effective signal to conspecifics, while other species might miss the miniscule motion.

Alternatively, a regular cone mosaic could improve chromatic resolution by uniformly distributing cones with differing spectral sensitivities across the retina, to help in adapting the visual system to the spectral composition of the photic environment through the relative sizes and ratios of individual cone types (Fernald, 1981; van der Meer, 1992). Accordingly, *T. delaisi*'s consistent mosaic could grant it acute color vision throughout its visual field. Additionally, the higher ratio of long-wavelength-sensitive twin cones in the foveal area (Fig. 8) increases sensitivity to the respective parts of the spectrum and suggests that light of longer wavelengths carries important information for *T. delaisi*, maybe helping to distinguish camouflaged prey, predators, or conspecifics from the background.

### Triple cones

The existence of different types of triple cones in the retinae of some teleost species is well-documented, but how they are distributed in the retina is often left out or mentioned only in passing (Hess, 2009). There is not a single triple cone map among the hundreds of entries in Clinical and Experimental Optometry's retinal topography maps database (Collin, 2008). Hence, this study is among the first to systematically analyze and map the triple cone distribution across an entire retina. *T. delaisi*'s triple cones were exclusively found in an area adjacent to the optic nerve head in the dorso-temporal retina (Fig. 8), which seems to be a common location for triple cones (Hess, 2009). Due to the localized occurrence, the total population was estimated to be only 5000 triple cones, or roughly one percent of all cones, which is similar to the proportion found in most other species (Hess, 2009). The exact function of triple cones is yet unclear. In some species they only occur after regeneration of previously mechanically damaged retina and are regarded as malformations (Cameron and Easter, 1995; Cameron and Powers, 2000). In other species, however, a role in vision seems evident, especially where triple cones dominate parts of or the entire retina (Hess, 2009). Intriguingly, Cameron's (1995) description of the triple cones' location, morphology, and arrangement after regeneration in the retina of the green sunfish matches the situation found in *T. delaisi* almost exactly. While this seems to suggest that the presence of triple cones in *T. delaisi* might also be the result of regenerative processes, it seems unlikely given the consistency of their occurrence and the naive condition of all study individuals. A visual function is more likely, because the triple cones are the dominant cone type where they occur, and the RGC density in the triple cone area is both relatively high and continuous with that of the surrounding retina. In some fishes, like the cutlips minnow and several anchovy species (Collin et al., 1996; Kondrashev et al., 2012), the triple cones consist of unequal members. In anchovies, it could further be shown that the lateral and central components exhibit different spectral sensitivities, which may suggest triplecones serve colour discrimination in the respective species (Kondrashev et al., 2012). In *T. delaisi*, however, the triple cones consist of morphologically equal members and, unfortunately, data on their spectral sensitivity is currently not available, as they were missed by Bitton et al. (2017) in their MSP study. Therefore, their role in the vision of *T. delaisi* remains unclear.

### Foveal structure

Many teleost retinae feature an area centralis, while foveae are less common (Collin and Pettigrew, 1988c; b; Nicol and Somiya, 1989; Collin, 2008). *T. delaisi*'s type of *fovea* with a steep, convexiclivate pit and incompletely displaced inner retinal layers is also found in syngnathids and the sandlance, who also share some aspects of their ecology with triplefins (Collin and Collin, 1988b; 1999). The curvature of the foveal pit combined with the higher refractive index of the retinal tissue compared to the vitreous create optical effects that may enhance multiple visual functions in the foveal area. One commonly suggested effect is the magnification of an image in the center of the fovea (Walls, 1937; Snyder and Miller, 1978). This effect is generated at the base of the foveal pit and requires it to have a smooth and symmetrical curvature.

*T. delaisi*'s foveal pit, however, can vary from a concentric dimple to a slit-like trough, which makes a consistent and precise optical magnification effect unlikely. Other studies suggest that image distortion by the foveal slope could inform about the focal state of the eye, help distinguish small targets from the background, and improve motion detection (Pumphrey, 1948; Harkness and Bennet-Clark, 1978; Steenstrup and Munk, 1980). These potential functions, which are not mutually exclusive, would be more plausible and clearly advantageous in *T. delaisi*'s ecological context. Unfortunately, none of the alleged functions has ever been tested experimentally, since the crucial factor of foveal slope might be impossible to manipulate. Accordingly, we too, can only speculate about its function in *T. delaisi*.

Another optical function could fall to the hemispherical thickening of the circumfoveal retina, especially if the foveal pit is very narrow or even closed. Inversely analogous to the telephoto lens effect in the steep fovea of raptors (Snyder and Miller, 1978), refraction at the convex interface between vitreous and retina could shorten the overall focal length of the eye and shift the triplefin's focal range further towards short distances. As a consequence, the area centralis might be unable to focus far away objects. Even if the high-acuity zone were inherently myopic, this may not be a disadvantage to *T. delaisi*, since distances greater than a few body lengths are probably irrelevant for small benthic fishes.

### Spatial resolving power

The major factors determining the spatial resolving power (SRP) of an eye are the number of neural elements in the retina and the quality of the optical system of the eye (Land and Nilsson, 2012). In terms of retinal elements, previous studies have often used either photoreceptor or RGC densities to estimate SRP, as reviewed by Caves et al. (2017). While the RGC are unquestionably the units of neural signaling between retina and the visual centers of the brain, using their density directly to estimate spatial resolution can be misleading, especially in species with a pronounced fovea. By definition, foveae are associated with a partial to complete displacement of RGCs, which causes a decreased RGC density in the foveal center, while the highest density is found circumfoveally. Since foveae are retinal specializations for high acuity, it is reasonable to assume that each receptor contributes an individual signal, *i.e*. the foveal summation rate is 1:1, at least in retinae adapted to photopic conditions. The displacement of RGC, however, can make the summation rate seem lower than it actually is and cause acuity to be underestimated. Our methodology did not allow us to estimate the actual summation rate in *T. delaisi*, but it is interesting to note that the RGC density stays relatively high throughout the retina, and that its average (32,000 cells mm^−2^) almost matches the average density of all cones (33,500 cells mm^−2^). Assuming that rods have high summation rates, as they serve scotopic vision, they would utilize only a fraction of the available RGCs. This would allow a relatively low summation rate for cones throughout the retina, and indicate that acuity is prioritized over sensitivity even in the peripheral retina.

Our photoreceptor-spacing-based estimates of *T. delaisi*'s anatomical SRP resulted in an acuity range of 6.7 - 9.0 cpd (9.0 - 6.7 arc minutes). This is supported by similar values reported for another triplefin species (Pankhurst et al., 1993) and also matches the overall average visual acuity of 8.4 ± 6.5 cpd found among 159 teleost species, as recalculated from the data in Caves et al. (2017). Based on Caves et al.'s methodology, *T. delaisi*'s acuity might be well above average for its "relative eye investment", a factor relating to the expected eye size for a given body size. Interestingly, the acuity of the smallest investigated species, as well as *T. delaisi*'s, is higher than would be expected for their sizes (Caves et al., 2017). This stresses once more the importance of acute vision in small fishes, as it can help to spot minute prey, cryptic conspecifics, intruders, or threats at greater distances, and reduces the required patrolling in their territory. A SRP of 6.7 - 9.0 cpd means *T. delaisi* could resolve a 1 mm target, like a copepod or small gammarid, at 38 - 52 cm (Fig. 7), which implies that it could monitor a large part of its ~1 - 3 m^2^ territory (Goncalves and Almada, 1998) from a stationary position, which is a typical component of their behavior.

Although *T. delaisi*'s retinal organization suggests a high investment in visual acuity, the receptor cell coverage in the center of the fovea, and thus the potential spatial resolution, is not maximized (Fig. 5D). The gaps in the mosaic pattern are due to the altered shape of the twin cones. The inner segment diameters of individual twin cone members and single cones in the fovea are reduced to ~2 μm, and their outer segments are approaching the physical size threshold beyond which they start losing part of the light energy and even interfere with their neighbors (Land and Nilsson, 2012). To maintain a maximal diameter, while the overall size is reduced, the twin cone members become rounded. As a consequence, the twin cones as a whole are more elongated, which skews the lucky-clover mosaic pattern, and decreases receptor coverage. That this pattern is nevertheless maintained in the fovea either suggests a developmental constraint, or that the benefit of a regular arrangement outweighs that of a maximized coverage. Maybe the fovea realizes a compromise between motion sensitivity, or maybe acute color vision, and absolute spatial resolving power.

## Conclusions

*T. delaisi*'s eye morphology, retinal organization and receptor composition show a host of adaptations that demonstrate the crucial role acute, color-and motion-sensitive vision plays in the life of a crypto-benthic micro-predator. The consistent and dominant presence of cones and the probably low summation ratio, as suggested by the high numbers of RGC even in the periphery, reveal a visual system adapted to a diurnal, photopic lifestyle in which acuity and color vision are prioritized over sensitivity. The elliptical eye shape and the antero-ventrally directed aphakic gap allow a greater range of lens movement and thus accommodation, which might be further boosted by the potential macro-lens effect of the hemispherical retinal thickening in the foveal region. The prominent *area centralis* with a convexiclivate fovea, and their dorso-temporal location in combination with the wide range of eye movements, allow high-resolution imaging of the substrate across a range of 120-130° in an antero-ventral direction. These complementary specializations are particularly advantageous for small benthic foragers, as they allow them to bring the substrate in their immediate vicinity into focus and scan it for potential prey at high resolution.

The consistent presence of all cone types, and their regular arrangement, possibly enhances color vision across the entire visual field. The change in the ratio of single to twin cones in the foveal region suggests a sensitivity boost for longer wavelengths, i.e. in the green, yellow, and red parts of the spectrum. All this might improve *T. delaisi*'s ability to distinguish and identify predators, prey, and conspecifics in its visually complex environment. An alternative, or additional, benefit of the regular arrangement of cones could be enhanced motion detection. The consistency of the mosaic throughout most of the retina might allow *T. delaisi* to notice minute movements in its entire visual field and draw its attention to the source, which it could then inspect with its fovea. The persistent maintenance of a regular mosaic even in the fovea, despite gaps in the receptor coverage, may indicate that the benefits to motion detection (or other functions) outweigh the gains of a maximized acuity. Finally, the extensive eye movement capabilities, the relatively high acuity even in the periphery, and the consistent motion and color sensitivity allow *T. delaisi* to monitor its entire surroundings, to assess threats or targets of interest “out of the corner of their eyes”, and to avoid moving their body and compromising their crypsis. All of these features make *T. delaisi* a prime example of how the visual systems of all species are adapted to their specific lifestyles and needs, and how much anatomical traits can tell us about a species' ecology. Despite a plethora of studies that have established the functions of most retinal features and their links to the respective species' ecology, some remain unresolved or are limited to speculation, like the function of triple cones and their peculiar, localized distribution in the retina of *T. delaisi*.

## Ethics statement

This study was carried out in accordance with German animal ethics legislation under notifications AZ. 13.06.2013 and AZ. 29.10.2014 issued to NKM by the animal welfare department of the district administration of Tübingen ("Regierungspräsidium Tübingen"). The protocol was approved by the animal ethics committee of the above authority.

## Authors' contributions

Conceptualization: RF and NM; Funding Acquisition: NM and SC; Resources: SC and NM; Methodology: RF, SC, and NM; Investigation: RF; Formal Analysis: RF; Writing – Original Draft: RF; Writing – Review & Editing: RF, SC, and NM; Supervision: SC and NM. All authors have read and approved the final manuscript.

The work presented in this study was funded by the German Science Foundation ("Reinhart Koselleck Project" grant Mi 482/13-1 to NM), the Volkswagen Foundation ("Experiment!" grant Az. 89 148 and 91 816 to NM), and by grants from the Australian Research Council (ARC) and the West Australian Government to SC. We are further thankful for the German Science Foundation's and the University of Tübingen Open Access Publishing Fund's support in making this study openly available.

## Acknowledgements

We would like to thank João Paulo Coimbra for his invaluable methodological involvement and critical input in the writing of this paper. We further thank Michael Archer, Tom Stewart, Jeremy Ullmann, and Gary Cowen for their assistance and guidance in producing the histological and MRI sections of *T. delaisi*’s head. We are deeply grateful to Radhika Puttagunta and the Hertie Institute for Clinical Brain Research, Tübingen, for allowing us access to their stereology set-up. Finally, we thank all colleagues and referees, whose critical comments helped improving this paper.

